# Natural Allelic Variations of Xenobiotic Enzymes Pleiotropically Affect Sexual Dimorphism in *Oryzias latipes*

**DOI:** 10.1101/000661

**Authors:** Takafumi Katsumura, Shoji Oda, Shigeki Nakagome, Tsunehiko Hanihara, Hiroshi Kataoka, Hiroshi Mitani, Shoji Kawamura, Hiroki Oota

## Summary

Sexual dimorphisms, which are phenotypic differences between males and females, are driven by sexual selection [1, 2]. Interestingly, sexually selected traits show geographic variations within species despite strong directional selective pressures [3, 4]. However, genetic factors that regulate varied sexual differences remain unknown. In this study, we show that polymorphisms in *cytochrome P450* (*CYP*) *1B1*, which encodes a xenobiotic-metabolising enzyme, are associated with local differences of sexual dimorphisms in the anal fin morphology of medaka fish (*Oryzias latipes*). High and low activity CYP1B1 alleles increased and decreased differences in anal fin sizes, respectively. Behavioural and phylogenetic analyses suggest maintenance of the high activity allele by sexual selection, whereas the low activity allele may have evolved by positive selection due to by-product effects of *CYP1B1*. The present data can elucidate evolutionary mechanisms behind genetic variations in sexual dimorphism and indicate pleiotropic effects of xenobiotic enzymes.

## Highlights

Xenobiotic enzyme CYP1B1 alleles cause genetic variation in sexually selected traits. The high enzyme activity allele has been maintained by sexual selection. By-product effects of CYP1B1 can cause reduced sexual dimorphism.

## Results and Discussion

Charles R. Darwin proposed that sexual selection drives sexual dimorphisms and speciation according to the reproductive strategy of each species [1]. Under such strong directional selection, alleles that enhance sexually selected traits are predicted to spread and fix within populations [2], leading to increased sexual dimorphisms and decreased genetic diversity [5] due to more successful mating of individuals with specific pronounced traits. However, phenotypic variations in the degree of sexual dimorphism are often found among wild populations within species. Evolutionary biologists have long discussed this paradox [3, 4], and several explanations have been proposed. Firstly, sexually selected traits can be indicative of a male’s quality and condition [6]. Under this ‘indicator model’, alleles that contribute to male qualities can influence sexually selected traits, resulting in corresponding phenotypic variations [7]. Secondly, ecology can influence the degree of sexual dimorphism [8], even in cases where sexual selection acts on male or female traits [9], leading to increased sexual differences and competitive mating that selects phenotypes that are advantageous for reproductive success but are disadvantageous for survival [10, 11]. Under this ‘trade-off model’ between sexual and natural selection, it is predicted that genes related to sexual dimorphism have pleiotropic functions, and may underpin differences in sexual dimorphisms as by-products of environmental adaptation. As a result, sexually selected traits can stabilise at an equilibrium point of both pressures, varying with environmental differences between local populations.

To unravel this paradox, we performed experiments using medaka fish (*Oryzias latipes*) that are prevalent throughout East Asia [12, 13] (Figure 1A) because they are an excellent animal model for studies on phenotypic variations [14]. Specifically, this model allows functional genetic comparisons among geographically local populations with abundant genetic diversity [15, 16], according to geographical differences in sexual dimorphisms of anal fin morphology [17, 18] (Figure 1B). Males from the Tanabe population (TN), which is a local population name, have larger anal fins, which are shaped as parallelograms and show clear sexual dimorphism (Figure 1C). In contrast, males from the Maegok population (MG) have smaller right-angled triangle-shaped anal fins showing little sexual dimorphism (Figure 1C). This anal fin trait is characterised by genetic architectures [17] and may have developed through competition, as indicated by the sexual function of the anal fin, which is used to hold females during mating (movie S1). Indeed, assuming that anal fin size variations affect mating behaviours, they may be positively correlated with reproductive success [19] and may reflect strong sexual selection. Thus, we determined whether males with larger anal fins had greater reproductive success by observing competitive mating behaviours of two males (from Tanabe and Maegok) and one female (from Tanabe or Maegok) medaka fish in a tank. Regardless of genetic background, the female mated more frequently with males carrying larger anal fins (Figure 1D; Spearman’s rank correlation, *rho* = 0.470, *p* = 0.004; Table S1), indicating that males with larger anal fins reproductively succeed those with smaller anal fins. Thus, anal fin morphology is significantly associated with reproductive behaviour and represents a feature of sexual selection in medaka fish. This geographical difference in medaka fish sexual dimorphism provides a unique opportunity to examine phenotypic variations that underlie sexually selected traits.

**Figure 1:**
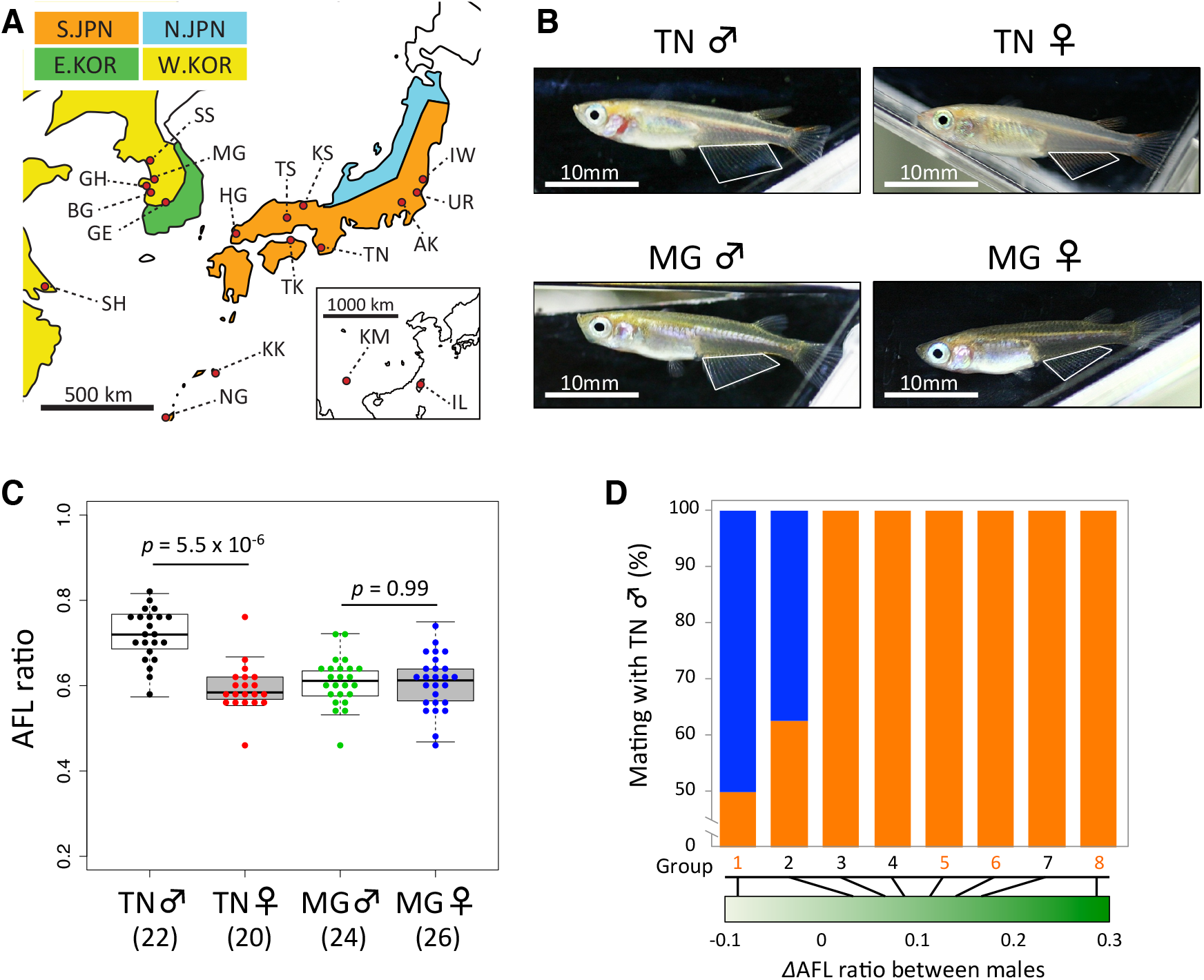
Phenotypic diversity among medaka populations. **(A)** Map of original medaka (*Oryzias latipes*) habitats; each colour represents geographically local populations. AK, HG, IW, KK, KS, NG, TK, TN, TS, UR, BG, GE, GH, IL, KM, MG, SH and SS are abbreviations for Akishima, Hagi, Iwaki, Kikai, Kasumi, Nago, Takamatsu, Tanabe, Tessei, Uridura, Bugang, Gwangeui, Guhang, Ilan, Kunming, Maegok, Shanghai and Samsan, respectively. Each color represents the location of *Oryzias latipes* populations based on mitochondrial genetic grouping; orange, Southern Japanese (S.JPN); cyan, Northern Japanese (N.JPN); green, Eastern Korean (E.KOR); yellow, Western Korean (W.KOR). **(B)** Differences in anal fin morphology between Tanabe and Maegok. **(C)** Sex-based variations in anal fin morphology between Tanabe and Maegok. Boxplots represent five-number summary statistics for each group, with lower and upper error bars indicating minimum and maximum observations, the tops and bottoms of boxes represent third and the first quartiles, respectively, and the middle bar represents the median. Numbers in parentheses represent numbers of individual fish. Significant differences were identified using Kruskal-Wallis’ test and Scheffe’s type multiple tests; \**p* < 0.05 (See Supplemental Experimental Procedures). **(D)** Frequencies of successful mating events with male fish from Tanabe (orange bars) are ordered by increasing differences (Δ) in AFL ratios between males (green gradient bar). Black and orange numbers represent Tanabe and Maegok females in the tank, respectively. See also Tables S1 and Movie S1.

Male anal fin sizes can be correlated with internal densities of estrogen in local medaka populations [17]. Sex steroid hormones such as estrogen often affect the development of sexual dimorphic traits [20], and are synthesised by various vertebrate cytochrome P450 (CYP) enzymes (KEGG PATHWAY: map00140). These enzymatic functions have been well studied because human CYPs play an important role in xenobiotic metabolism and drug digestion [21]. Our preliminary screening studies showed high degrees of polymorphism among medaka *CYPs* and suggested that differing activities of *CYP1A* and *CYP1B1* alleles contribute to differential regulation of sex steroid metabolism between medaka populations from various locations. Indeed, *CYP1A* and *CYP1B1* were highly polymorphic and differentiated across medaka populations, and were also involved in estrogen metabolism in fish [22] (Figure S1 and Supplemental Experimental Procedures). Therefore, we investigated the role of *CYP* polymorphisms and the ensuing molecular mechanisms that lead to geographical variations in medaka anal fin morphology, and elucidated the evolution of sexual dimorphisms under *CYP* genes. To this end we (1) measured differences in enzyme activity between CYP alleles using biochemical assays, (2) examined whether these alleles were associated with differences in degrees of sexual dimorphism and (3) identified selective pressures acting on each allele of *CYP1B1* by identifying mutations acquired from common ancestors.

Assuming that associations exist between *CYP1A* and *CYP1B1* variants and medaka anal fin morphology, functional allelic differences may exist between the two populations. To assess variations in CYP enzyme activities, we cloned the first and second most frequent haplotypes of *CYP1A* and *CYP1B1* from S.JPN (Southern Japanese group) and W.KOR (Western Korean group), respectively, into pUAST expression vectors and expressed these in *Drosophila* S2 cells. Because medaka from Tanabe carried major *CYP* haplotypes (H02 in oCYP1A and H03 in oCYP1B1; Figure S1), enzyme activities were expressed relative to those of CYP1A and CYP1B1 in fish from Tanabe. Kasumi’s CYP1A haplotype (H02 oCYP1A) exhibited the highest relative activity (1.24-fold that from Tanabe medaka, *p* = 0.019), but large differences were not observed between the four haplotypes examined (range, 0.70–1.24; Figure 2A). In contrast, relative enzyme activities of CYP1B1 haplotypes were significantly lower than those of the Tanabe haplotypes (Figure 2B). Maegok haplotypes (H08 oCYP1B1) had substantially lower activities (26%; *p* = 0.017) than the major Tanabe haplotype (H03 oCYP1B1). Subsequent western blotting experiments confirmed that the observed inter-haplotype differences in CYP1B1 activity were not due to relative protein expression, but due to differences in amino acid sequences (Figure S1F). Thus, the two CYP1B1 haplotypes H03 (Tanabe-type) and H08 (Maegok-type) had significantly higher and lower enzyme activities, respectively.

**Figure 2:**
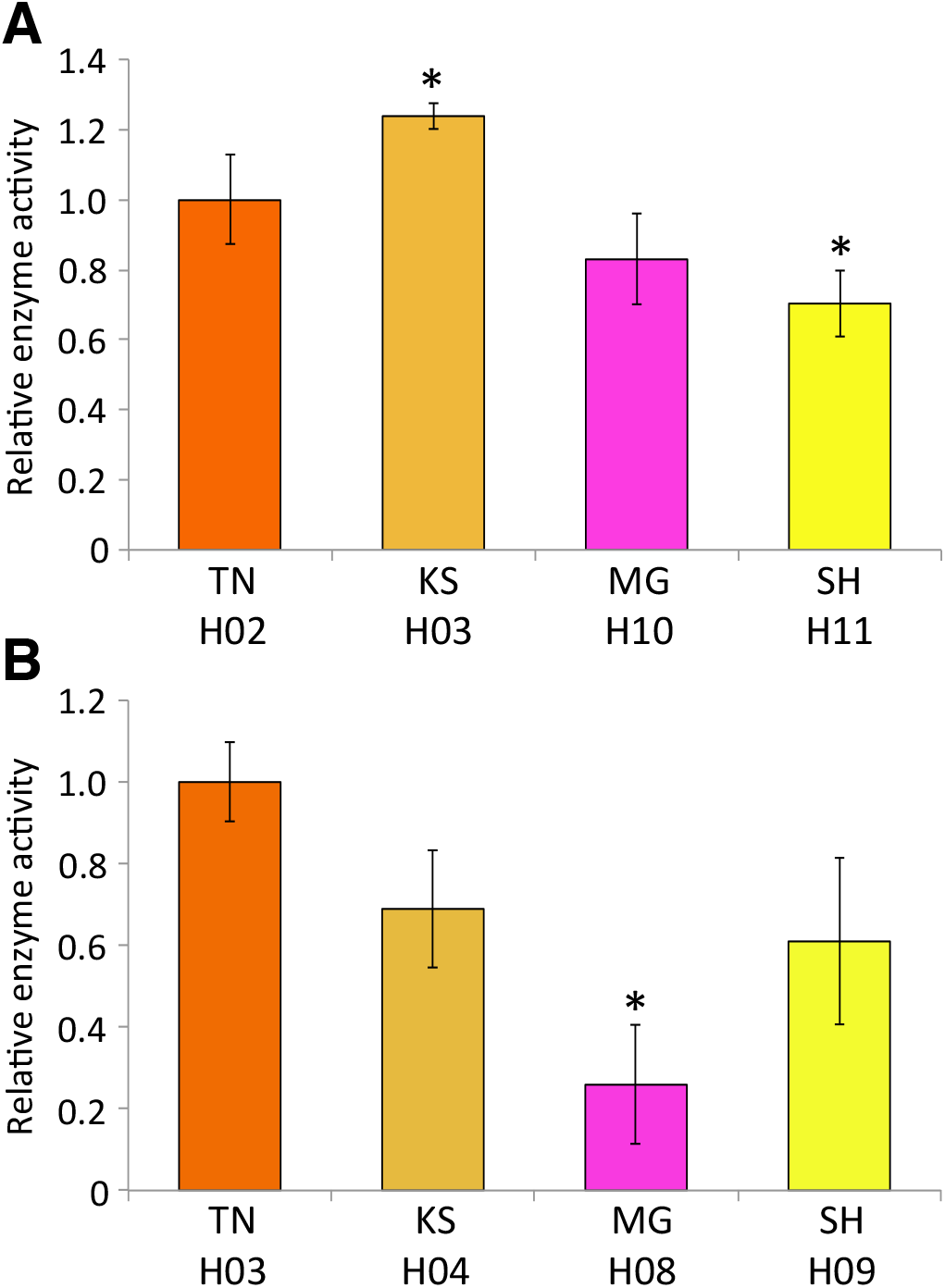
Relative enzyme activities of medaka CYP1A and CYP1B1. The x-axis displays the names of local wild medaka populations and corresponding CYP1A (**A**) and CYP1B1 (**B**) haplotypes. Each bar represents the mean ± S.D. from multiple independent samples. Each colour corresponds with that in Figure S1A-D. Significant differences were identified using Games–Howell tests; \**p* < 0.05.

We used genetic and phenotypic data from 102 F2 fish obtained by crossing Tanabe and Maegok medaka fish to examine the relationship between *CYP1B1* genotype and anal fin morphology. AFL ratios (A-AFL/P-AFL; see Supplemental Experimental Procedures) significantly differed between the sexes in *CYP1B1* Tanabe-type homozygotes (TT) and in heterozygotes of Tanabe and Maegok types (TM), but did not in Maegok-type homozygotes (MM; *p* = 0.027, *p* = 2.7 × 10^−4^ and *p* = 0.22, respectively; Figure 3A). Subsequently, we calculated *C* score-based Mahalanobis’ generalised distances (*D*^2^), which were corrected for the influence of differences in the sizes of analysed traits between groups of F2 individuals that were categorised according to *CYP1B1* genotypes (Table S2). *D*^2^ values indicated pronounced sex differences, and between-sex *D*^2^ values among Tanabe-type homozygotes (TT♂–TT♀) were approximately 1.6 times higher than those among Maegok-type homozygotes (MM♂–MM♀). These were presented in 2D deployments of *D*^2^ values using classical multidimensional scaling (cMDS; Figure 3B). TT males and females were more distant from each other than MM and TM males and females, indicating that homozygote males and females from Tanabe were more sexually dimorphic than those from Maegok. Thus, these results show that *CYP1B1* is a predominant predictor gene for sexually selected traits.

**Figure 3:**
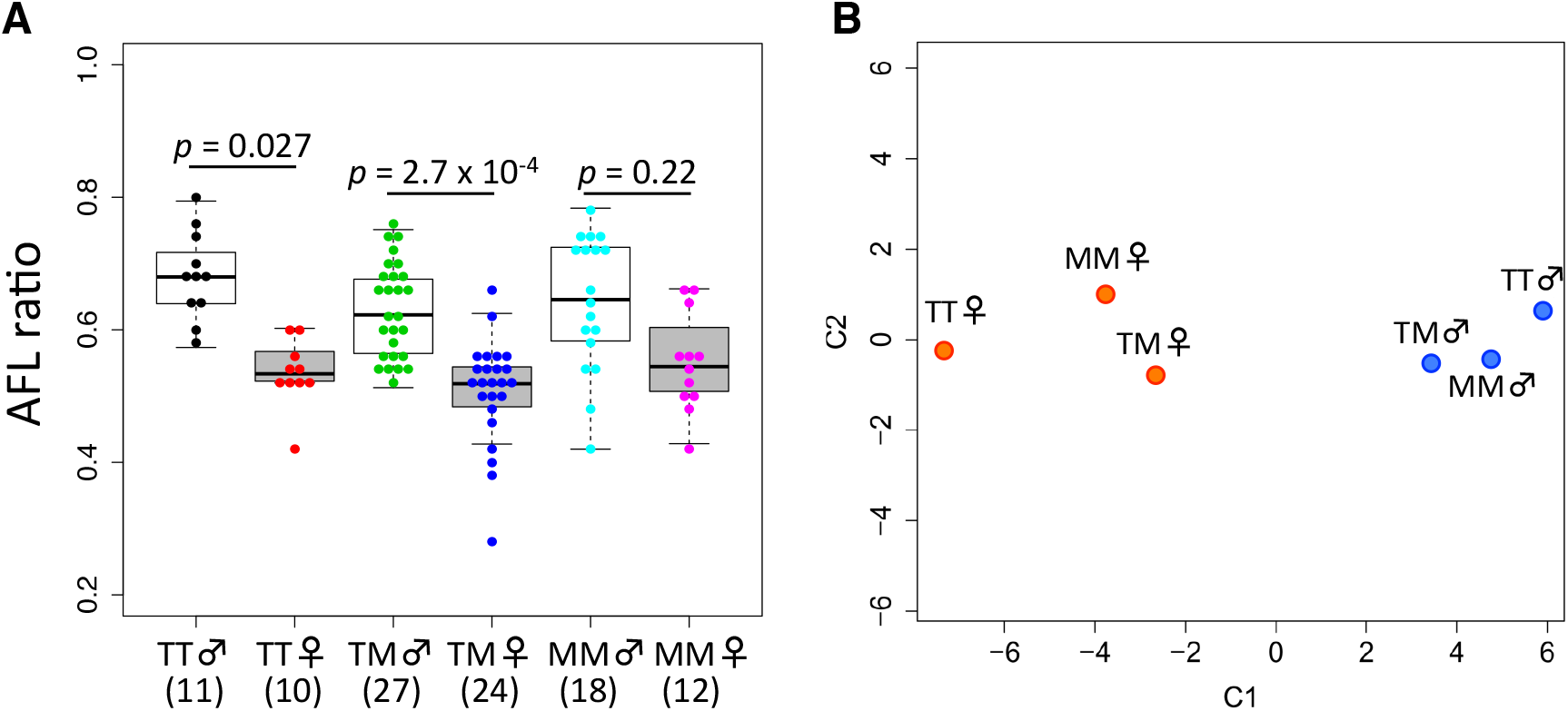
Comparisons of anal fin morphology among F2 individuals. **(A)** Boxplots show differences in AFL ratios between combinations of *CYP1B1* genotypes and sexes. Coloured dot plots show distributions of data within each group; TT, homozygote from TN; TM, heterozygote from TN and MG; MM, homozygotes from MG. Significant differences were identified using Kruskal-Wallis’ test and Scheffe’s type multiple tests; \**p* < 0.05 (See Supplemental Experimental Procedures). **(B)** Multidimensional scaling plots of *C* score-based Mahalanobis’ generalised distances (*D*^2^) show marked morphological sex differences between *CYP1B1* genotypes (See also Table S2). Blue and red dots represent males and females, respectively.

Because males with larger anal fins mated more successfully (Figure 1D), we expected that anal fin-derived sexual selection would be accompanied by alleles that give rise to high enzyme activity. To determine selective pressures on *CYP1B1*, we estimated likelihood ratios [23], tested constancy of non-synonymous SNPs over synonymous SNPs ratios (dN/dS = ω) among evolutionary lineages and showed significant non-constancy of the ω value for each lineage (*p* < 0.05; Figure S2A). Interestingly, only a single amino acid substitution was found in the lineage from the common ancestral allele compared with the Tanabe-type allele, as indicated by a ω value (0.001) of significantly less than 1 (Figures 4A and S2A). In the S.JPN, nucleotide diversity of the *CYP1B1* coding region was lower than that of other *CYP* genes (Figure 4B), indicating evolutionary conservation of *CYP1B1*. These observations suggest that purifying selection (background selection) has operated on the Tanabe-type allele. Moreover, reconstructed ancestral CYP1B1 showed high enzyme activity equivalent to that of the Tanabe-type allele (Figure S2B), suggesting functional conservation of this allele for the 5 Myr since divergence from the common ancestral population (Table S3). These results support the association between the high enzyme activity (Tanabe-type) allele and the preponderance of sexual dimorphism, and strongly suggest that this allele has been maintained by sexual selection.

**Figure 4:**
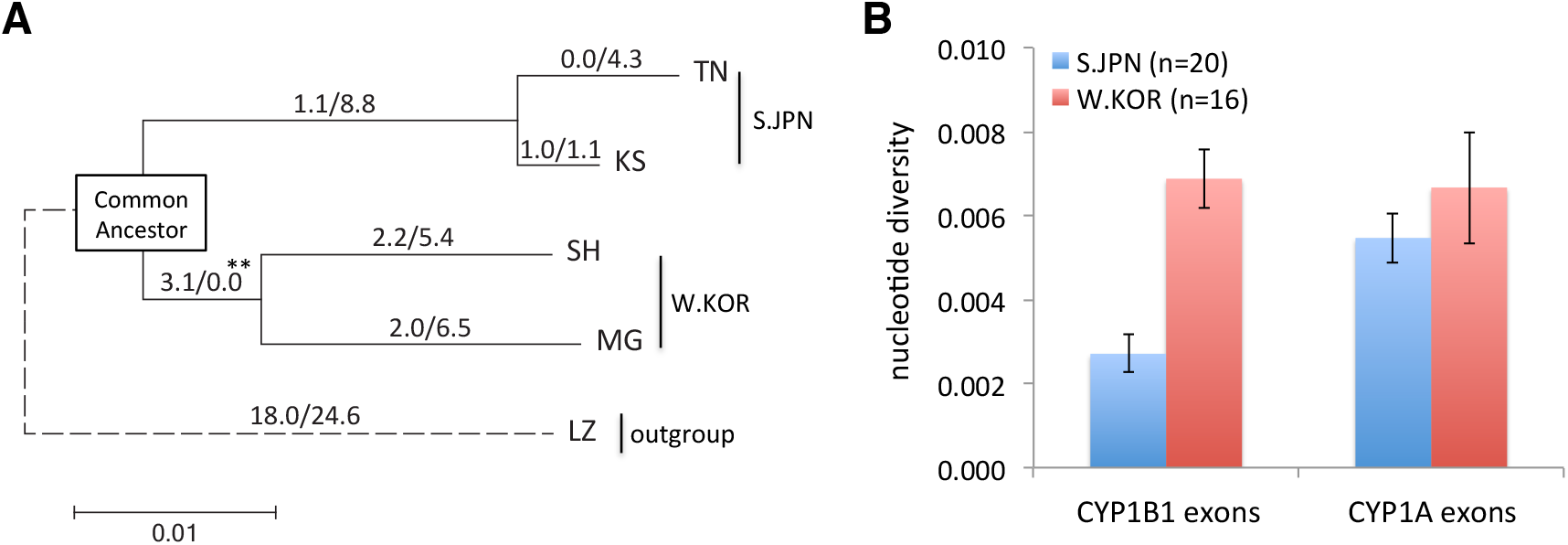
Genetic diversity of *CYP1B1*. **(A)** Maximum likelihood tree with synonymous and non-synonymous SNPs of *Oryzias* genus. LZ (*O. lusonensis*) from the Philippines was used as the out-group. The square on the tree indicates common ancestor of S.JPN and W.KOR; significant likelihood ratio test, \*\**p* < 0.02 (See Figure S2 and Supplemental Experimental Procedures). **(B)** Comparisons of nucleotide diversity calculated on nucleotide sequences of *CYP* exons; each bar represents the mean ± S.D., n indicates the number of chromosomes.

Because Maegok females mate more readily with males carrying larger anal fins (Figure 1D), the ‘indicator model’ was ruled out, and an additional process to sexual selection may have contributed to reduced sexual dimorphism. In the phylogenetic tree, the w value (3.1/0.0) significantly increased only on the branch of the ancestral allele that was common to both Maegok and Shanghai alleles (*p* < 0.02; Figures 4A and S2A), implying that positive selection operated on this allele. Given that previous studies have shown toxicity of CYP1B1 metabolites of 17β-estradiol [24], and survival advantages of CYP inactivation in vertebrates under certain conditions [21, 25], selective pressures on the common ancestral population may have been environmental (e.g. water pollution). Hence, positive selection on the ancestral branch indicates that the low activity allele may have been advantageous for survival of the ancestral population. These results indicate that adaptive evolution of this allele may have reduced sexual dimorphism as a by-product. Indeed, the effects of positive selection may have predominated over those of sexual selection.

These phenomena in medaka *CYP1B1* may have convergently evolved in other vertebrates, as indicated by pigmentation in sticklebacks and humans [26]. The human polymorphic variants CYP1B1*1, CYP1B1*2, CYP1B1*3 and CYP1B1*4 have differing enzyme activities [27]. Among these, CYP1B1*3 exhibits the highest enzyme activity and the highest frequency among Africans and Europeans (Figure S3A). Interestingly, CYP1B1*1 had lower activity than CYP1B1*3 but was the most prevalent haplotype in East Asians. A study of morphological anthropology shows that sex differences in tooth crown sizes are 2-fold larger in Africans and Europeans than in East Asians [28] (Figure S3B) and are coincident with the frequency distribution of the *CYP1B1*3* haplotype. This pattern is similar to the present results from medaka fish, suggesting potential analogy to the evolution of sexual dimorphism in humans. Hence, examinations of sex-difference diversities among humans could be considered in terms of the competing effects of sexual and natural selection of *CYP1B1*.

This study shows that polymorphisms in genes encoding xenobiotic metabolising enzymes can decrease sexual dimorphism under adaptive evolution, despite the strong phenotypic influence of sexual selection. This new mechanism is distinct from the widely accepted mechanism that all sexually dimorphic traits are fundamental products of sex-limited gene expression [20]. Although whole genome analyses are required to identify genes related to the differences between local populations, our candidate gene approach considerably examined differences in traits between geographically isolated populations of vertebrates. In addition, this study indicates analogous relationships between polymorphisms and phenotypic traits in wild medaka populations to those in humans. Comparisons of distinct vertebrate populations with common genetic polymorphisms may provide insights into the evolutionary significance of functional differences between alleles.

## Accession Numbers

Nucleotide sequence data were deposited into the international DNA database DDBJ/EMBL/GeneBank (accession numbers: AB829613–AB829721 and AB830922– AB830964).

## Acknowledgements

We thank Mrs. Shizuko Chiba, Mrs. Sumiko Tomizuka, Dr. Atsuko Shimada and Professor Emeritus Akihiro Shima (University of Tokyo) for keeping medaka stocks from wild populations. We thank Dr. Naoki Osada (National Institute of Genetics) for his useful comments, Takeshi Matsuya and Tetsuya Yamada for their support in western blotting experiments and measurements of CYP enzyme activity. We are also thankful to the National BioResource Project (NBRP)-medaka for providing wild medaka. This study was supported in part by Grants-in-Aid for Challenging Exploratory Research from the Japan Society for the Promotion of Science, No. 21657065 and No. 23657164 to H.O. and S.K, respectively. T.K. was supported by a Grant-in-Aid from JSPS Research Fellow (22-6207).

